# A novel yeast hybrid modeling framework integrating Boolean and enzyme-constrained networks enables exploration of the interplay between signaling and metabolism

**DOI:** 10.1101/2020.09.11.290817

**Authors:** Linnea Österberg, Iván Domenzain, Julia Münch, Jens Nielsen, Stefan Hohmann, Marija Cvijovic

## Abstract

The interplay between nutrient-induced signaling and metabolism plays an important role in maintaining homeostasis and its malfunction has been implicated in many different human diseases such as obesity, type 2 diabetes, cancer and neurological disorders. Therefore, unravelling the role of nutrients as signaling molecules and metabolites as well as their interconnectivity may provide a deeper understanding of how these conditions occur. Both signalling and metabolism have been extensively studied using various systems biology approaches. However, they are mainly studied individually and in addition current models lack both the complexity of the dynamics and the effects of the crosstalk in the signaling system. To gain a better understanding of the interconnectivity between nutrient signaling and metabolism, we developed a hybrid model, combining Boolean model, describing the signalling layer and the enzyme constraint model accounting for metabolism using a regulatory network as a link. The model was capable of reproducing the regulatory effects that are associated with the Crabtree effect and glucose repression. We show that using this methodology one can investigat intrinsically different systems, such as signaling and metabolism, in the same model and gain insight into how the interplay between them can have non-trivial effects by showing a connection between Snf1 signaling and chronological lifespan by the regulation of NDE and NDI usage in respiring conditions. In addition, the model showed that during fermentation, enzyme utilization is the more important factor governing the protein allocation, while in low glucose conditions robustness and control is prioritized.

**Author summary:** Elucidating the complex relationship between nutrient-induced signaling and metabolism represents a key in understanding the onset of many different human diseases like obesity, type 3 diabetes, cancer and many neurological disorders. In this work we proposed a hybrid modeling approach, combining Boolean representation of singaling pathways, like Snf11, TORC1 and PKA with the enzyme constrained model of metabolism linking them via the regulatory network. This allowed us to improve individual model predictions and elucidate how single components in the dynamic signaling layer affect the steady-state metabolism. The model has been tested under respiration and fermentation, reveling novel connections and further reproducing the regulatory effects that are associated with the Crabtree effect and glucose repression. Finally, we show a connection between Snf1 signaling and chronological lifespan by the regulation of NDE and NDI usage in respiring conditions.

## Introduction

Biological systems are of complex nature comprising numerous dynamical processes and networks on different functional, spatial and temporal levels, while being highly interconnected (Walpole et al., 2013). The field of systems biology faces the great challenge of elucidating how these interconnected systems work both separately and together to prime organisms for survival. One of such phenomena is the cells ability to sense and respond to environmental conditions such as nutrient availability. In order to coordinate cellular metabolism and strategize, the cell needs an exact perception of the dynamics of intra- and extra-cellular metabolites (Y. P. Wang & Lei, 2018). Simultaneously, nutrient-induced signaling plays a pivotal role in numerous human diseases like obesity, type 2 diabetes, cancer and ageing (Coughlan et al., 2014; Weidong Li et al., 2015; Salminen & Kaarniranta, 2012; Steinberg & Kemp, 2009). Therefore, unravelling the role of nutrients as signaling molecules and metabolites as well as their interconnectivity may provide a deeper understanding of how these conditions occur.

Yeast has long been used as a model organism for studying nutrient-induced signaling (Conrad et al., 2014a). Two major classes of nutrients include carbon and nitrogen. Carbon-induced signaling acts mainly through the PKA and SNF1 pathway while nitrogen-induced signaling acts through the mTOR pathway. The PKA pathway plays a major role in regulating growth by inducing ribosome biogenesis genes and inhibiting stress response genes(Broach, 2012). The SNF1 pathway is mainly active in low glucose conditions where it promotes respiratory metabolism, glycogen accumulation, gluconeogenesis and utilization of alternative carbon sources but it also controls cellular developmental processes such as meiosis and ageing (Ashrafi et al., 2000; Conrad et al., 2014a; Hedbacker & Carlson, 2008). The strongly conserved TORC1 pathway plays a crucial role in promoting anabolic processes and cell growth in response to nitrogen availability (Broach, 2012). Active TORC1 induces ribosomal protein and ribosome biogenesis gene expression (Lempiäinen et al., 2009; Marion et al., 2004) and represses transcription of genes containing STR and PDS elements in their promoter region (Lempiäinen et al., 2009). Despite the fact that Snf1, TORC1 and PKA pathways belong to the most well-studied pathways (Y. P. Wang & Lei, 2018), still, there is a lack of understanding both in the dynamics and the interactions leading to change in gene expression. It has been shown that glucose signaling is related to metabolism however the nature of this relationship remains unknown (Welkenhuysen et al., 2017). Numerous crosstalk mechanisms between these pathways have been described (Shashkova et al., 2015) and depending on their activity, they may influence the overall effect of the signaling process and thus the interaction with the metabolism (Welkenhuysen et al., 2019). In order to better understand the impact of cell signaling on metabolism, a systems biology approach is often implemented (Nielsen, 2017).

Typically, Boolean models have been developed to study the crosstalk between the Snf1 pathway and the Snf3/Rgt2 pathway (Christensen et al., 2009) as well as the Snf1, cAMP-PKA, and Rgt2/Snf3 pathways (Welkenhuysen et al., 2019). In mammalian cells, Boolean models have been used to evaluate the conflicting hypothesis of the regulation of the mTOR pathway (Sulaimanov et al., 2017) and to study crosstalk between mTOR and MAPK signaling pathways (Siegle et al., 2018). Since, signaling systems are not always strictly Boolean in its nature, where location, combinations of post-translational modifications as well as other interaction plays a role, different Boolean frameworks for handling these complex interactions have been developed (Romers et al., 2020; Welkenhuysen et al., 2019). In contrast, metabolism, also in itself a complex process, is often studied using Flux Balance Analysis (FBA), that enables prediction of biochemical reaction fluxes, cellular growth on different environments and gene essentiality even for genome-scale metabolic models (GEMs) (Lu et al., 2019; Monk et al., 2017; Yilmaz & Walhout, 2017). A major limitation of the use of GEMs together with FBA is the high variability of flux distributions for a given cellular objective (Mahadevan & Schilling, 2003), as FBA solves largely underdetermined linear systems through optimization methods. To overcome this problem, experimentally measured exchange fluxes (uptake of nutrients and secretion of byproducts) are incorporated as numerical constraints, however, such measurements are not always available for a wide variety of organisms and growth conditions.

The concept of enzyme capacity constraints has been incorporated into FBA in order to reduce the phenotypic solution space (i.e. exclusion of flux distributions that are not biologically meaningful) and diminish its dependency on condition-dependent exchange fluxes datasets (Adadi et al., 2012; Beg et al., 2007; Bekiaris & Klamt, 2020; Massaiu et al., 2019; Nilsson & Nielsen, 2016; Sánchez et al., 2017).

Notably, a method to account for enzyme constraints, genome-scale models using kinetics and omics (GECKO) (Sánchez et al., 2017)has been developed. GECKO incorporates constraints on metabolic fluxes given by the maximum activity of enzymes, which are also constrained by a limited pool of protein in the cell. This method has refined predictions for growth on diverse environments, cellular response to genetic perturbations, and even predicted the Crabtree effect in *S. cerevisiae*’s metabolism, but also proven to be a helpful tool for probing protein allocation and enabled the integration of condition-dependent absolute proteomics data into metabolic networks (Massaiu et al., 2019; Sánchez et al., 2017).

Following the holistic view of systems biology, hybrid models allow us to take the next step and combine many different formalisms to study the interconnectivity and crosstalk spanning different scales and/or systems. For example, to quantify the contribution of the regulatory constraints of an *E*.*coli* genome-scale model, a steady-state regulatory flux balance analysis (SR-FBA) has been developed (Shlomi et al., 2007), further the diauxic shift in *S. cerevisiae* has been studied by CoRegFlux workflow integrating metabolic models and gene regulatory networks(Banos et al., 2017). To bypass the need for kinetic parameters, a FlexFlux tool has been developed where metabolic flux analyses using FBA have been constrained with steady-state values resulting from the regulatory network(Marmiesse et al., 2015). This strategy has also been used in a hybrid model of the *Mycobacterium tuberculosis* where the gene regulatory network was used to constrain the metabolic model to study the adaptation to the intra-host hypoxic environment(Bose et al., 2018). However, to further study the impact of signaling on the metabolism, the complexity of the signaling systems itself and the crosstalk between interacting pathways need to be represented in a coherent manner.

To better understand the complex relationship between metabolism and signaling pathways, we created a hybrid model consisting of a Boolean module integrating the PKA, TORC1 and the Snf1 pathways as well as the known crosstalk between them and an enzyme-constrained module of *S. cerevisiae’s* central carbon and energy metabolism (Figure 1). The backbone of the presented model is a framework for utilizing the complex Boolean representation of large-scale signaling systems to further constrain an enzyme-constrained model (ecModel), where the activity of the transcription factors resulting from the Boolean module was used to constrain an ecModel of the central carbon metabolism. The proposed hybrid model is capable of reproducing the regulatory effects that are associated with the Crabtree effect and glucose repression and have further showed a connection between glucose signaling and chronological lifespan by the regulation of NDE and NDI usage in respiring conditions. In addition, the model showed that during fermentation, enzyme utilization is the more important factor governing the protein allocation, while in low glucose conditions robustness and control is prioritized.

**Figure 1.**
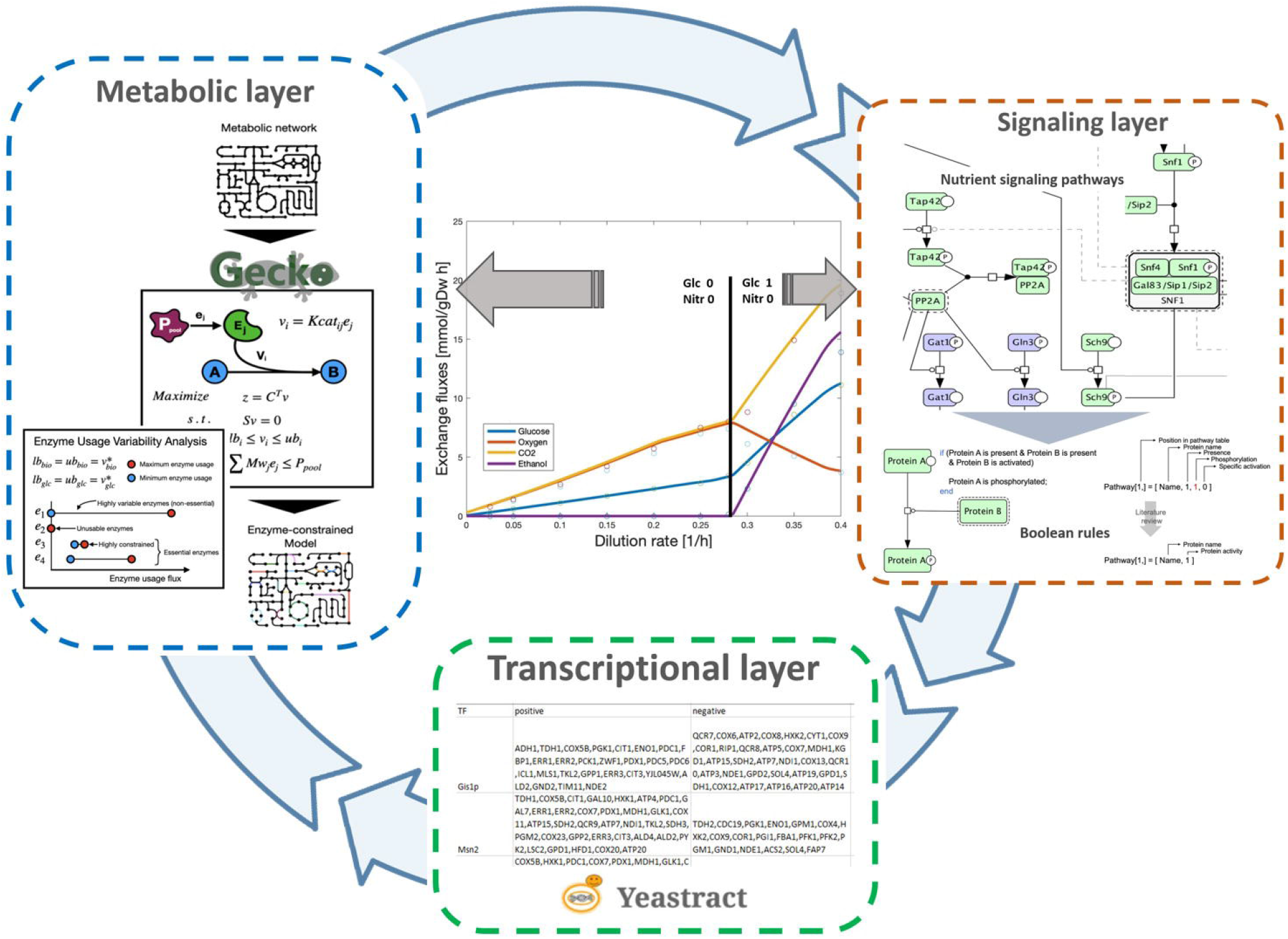
Schematic representation of the hybrid model. The hybrid model consists of a vector based Boolean module of nutrient signaling and a enzyme constrained module of the central carbon metabolism. The Boolean module is a dynamic module including Snf1, PKA and TORC1 pathway as well as crosstalk between them. The dynamic module reaches steady state and the activity of the transcription factors acts as input in a regulatory network constraining the enzyme constraint model of the central carbon metabolism. The solution is used to determine the activity of the Boolean input.

## Results

### Implemented Boolean signaling network is able to reproduce the general dynamics caused by glucose and nitrogen addition to starved cells

To verify the constructed Boolean model of nutrient-induced signaling pathways (Figure 2), cells were simulated from nitrogen and glucose starved conditions to nutrient rich conditions. The PKA pathway was activated upon glucose abundance via the small G proteins Ras and Gpa2. These proteins, in turn, activated the adenylate cyclase (AC) that induced processes leading to the activation of the catalytic subunit of PKA. Active PKA phosphorylated and therefore inactivated Rim15, thus the transcription factors Gis1, Msn2 and Msn4 became inactive.

**Figure 2.**
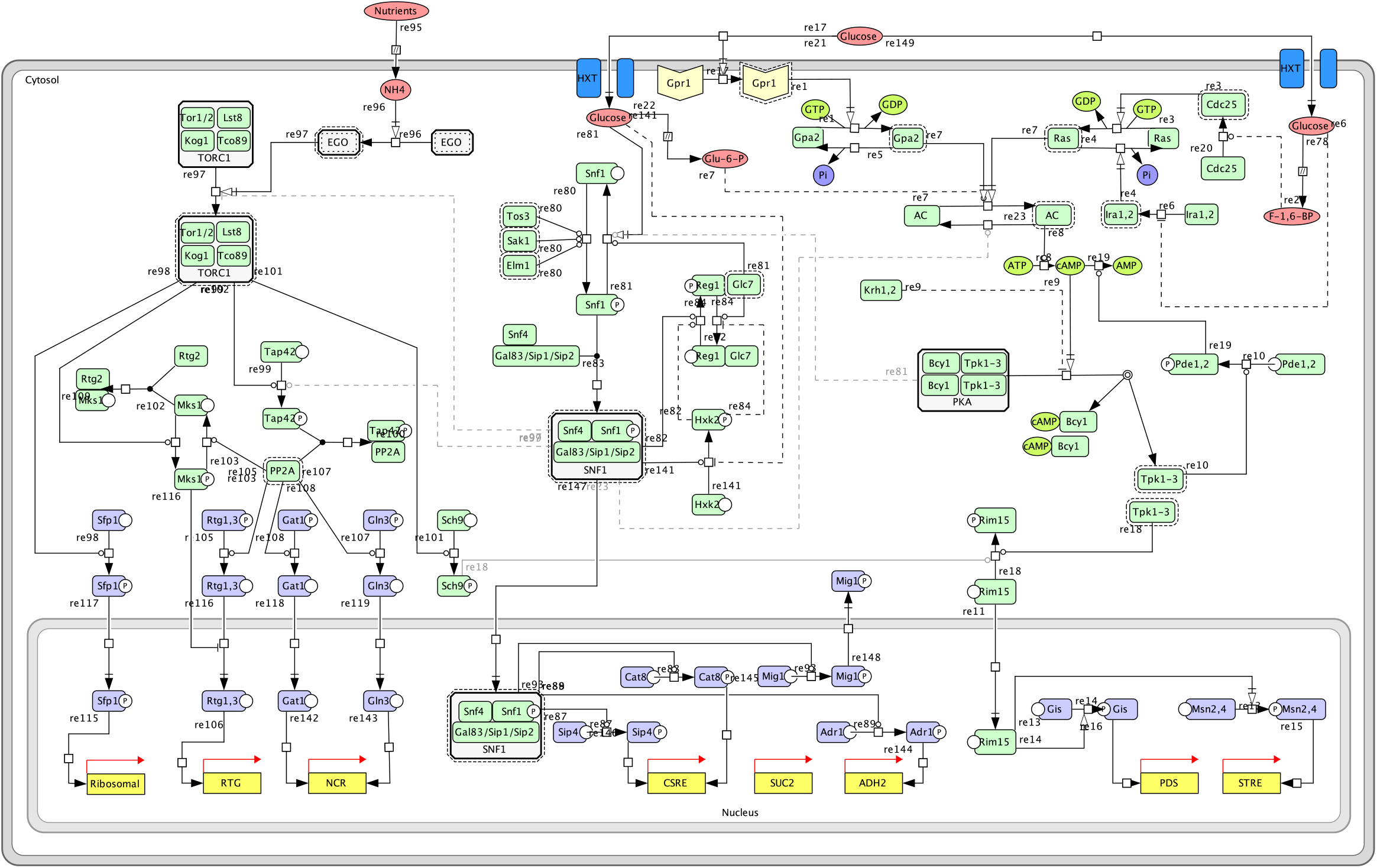
The Bolean module is a dynamic module including Snf1, PKA and TOR pathway as well as crosstalk between them. Crosstalk events between the pathways are depicted in grey. Unknown mechanisms are represented by dashed lines.

The SNF1 pathway is active when glucose is limited, while the addition of glucose causes Snf1 inactivation resulting in the activation of the transcriptional repressor Mig1 and the deactivation of Adr1, Cat8 and Sip4. However, the inactivation of Adr1 happened prior to Snf1 inactivation. This is due to the implemented crosstalk with the PKA pathway, where activated PKA inhibits Adr1 activity (Cherry et al., 1989).

Nutrient availability activates the TOR complex 1 which in turn phosphorylates Sch9 and Sfp1 resulting in the repression of Rim15 phosphorylation and the expression of ribosomal genes respectively. No change was observed in the activity of PP2A-regulated transcription factors Rtg1, Rtg3, Gat2 and Gln2. However, during the 8^th^ iteration, PP2A was active. In addition, Sch9 was not the main regulator of Rim15 activity in our simulations since PKA was activated prior to Sch9 and acted independently to regulate Rim15, either due to a gap in the model or a lack of complexity in our understanding of the signalling system (Supplementary Information S1).

### The Boolean model reveals interconnectivity and knowledge gaps in nutrient signaling pathways

To further investigate the impact of crosstalk in the Boolean model, knockouts of main components of each pathway (Snf1, Reg1, Tpk1-3 and Tor1,2) were simulated and compared to the wildtype in glc|nitr = 1|1 and glc|nitr = 0|0, see Figure S2. In nutrient-depleted conditions, only the Snf1 knockout had a significant impact. In the Snf1 pathway, Snf1 knockout affected all downstream targets leading to a transcription factor activity pattern that is usually observed in wildtype strains when glucose is available (Conrad et al., 2014b). It has been previously described that the phenotype of Snf1 mutants resembles the phenotype observed when the cAMP/PKA pathway is over-activated (Thompson-Jaeger et al., 1991). Although activation of the adenylate cyclase (AC) could be observed in the simulated knockout, PKA and the downstream targets were inactive due to the activity of the Krh proteins that inhibit PKA if no glucose is present in the Boolean model (T. Peeters et al., 2006). The Snf1 mutant showed defects in the TOR pathway upon glucose depletion leading to the activation of the PP2A phosphatase. The resulting activation of NCR and RTG genes and deactivation of ribosomal genes correspond to the phenotype one would expect if glucose but not nitrogen is available (Hughes Hallett et al., 2014) thus stressing the role of Snf1 in imparting the glucose state to the other nutrient-signaling pathways.

Under high nutrient availability, the Reg1 knockout showed almost the same effect on the SNF1 and TORC1 pathway as nutrient depletion. Only Adr1 activity was not affected which opposes the observations by Dombek and colleagues that described constitutive ADH2 expression in Reg1 mutant cells (Dombek et al., 1999).

An almost similar effect on the SNF1 and TORC1 pathways could be observed when Tpk1-3 knockout was simulated. This redundant effect was expected since impaired PKA activity was described to be associated with increased SNF1 activity(Barrett et al., 2012). Nevertheless, PKA knockout additionally induced Adr1 activation when SNF1-mediated activation could no longer be inhibited by PKA. The PKA knockout simulation showed strong effects on all three simulated pathways and may explain why strains lacking all three Tpk isoenzymes are inviable (Robertson & Fink, 1998).

The effects of Tor1 and 2 knockouts only affected the TORC1 signaling pathway. The simulated phenotype equalled the phenotype that is expected upon nitrogen depletion and glucose abundance and was therefore similar to the phenotype observed when simulating the Snf1 knockout in nutrient-starved cells. Besides, experimental observations revealed that impairing Tor1 and 2 function results in growth arrest in the early G1 phase of the cell cycle, as well as inhibition of translation initiation which are characteristics of nutrient, depleted cells entering stationary phase (Barbet et al., 1996). The fact that inactivation of TORC1 results in the inactivation of Sfp1 that regulates the expression of genes required for ribosomal biogenesis could be an indicator of this observation; however other TORC1-associated signaling mechanisms inducing translation initiation may likely be involved (Barbet et al., 1996).

### The hybrid model improves predictions of individual proteins and reveals a connection between regulation and chronological ageing

To verify the ecModel performance, the predictions of exchange reaction fluxes at increasing dilution rates were compared against experimental data (Van Hoek et al., 1998) (Figure S3), predictions showed a median relative error of 9.82% in the whole range of dilution rates from 0 to 0.4 h^-1^, spanning both respiratory and fermentative metabolic regimes. The hybrid model, including regulation, was further compared with the ecModel in their ability to predict protein abundances by comparing the predicted abundances to proteomics data from the literature in both respiratory and fermentative conditions (Doughty et al., 2020; Paulo et al., 2016). By adding the regulation layer, the prediction accuracy of individual protein abundances was drastically improved, reducing the mean absolute log_10_-transformed ratio between predicted and measured values (**r**) from 2.62 to 1.55 for respiration and from 3.56 to 2.32 for fermentation (Figure 3), which represents an average improvement in protein predictions by more than one order of magnitude for both conditions. Moreover, two sample Kolmogorov-Smirnov tests did not show significant statistical differences between model predictions and the available proteomics datasets. This large improvement is predominantly resulting from the utilization of more than one isoform for some reactions in the hybrid model in contrast to a pure ecModel, in which just the most efficient enzyme for a given reaction is used.

**Figure 3.**
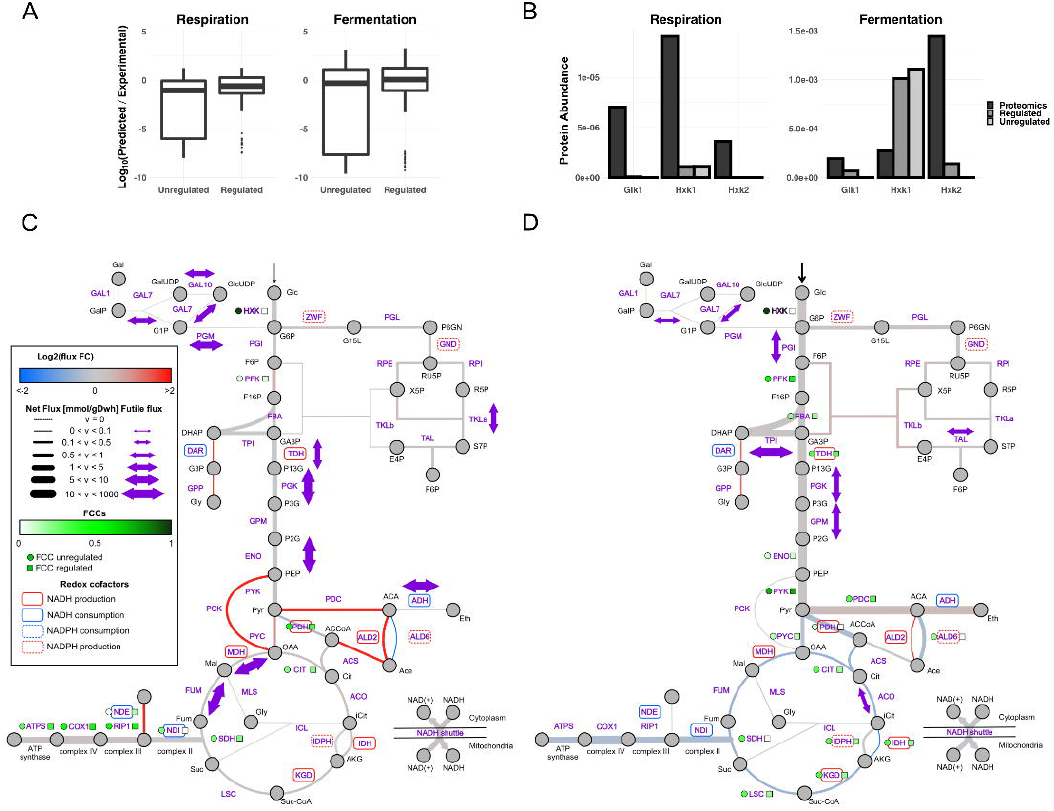
(A) Absolute log_10_-transformed ratio between predicted and measured values in respiration and fermentation. The plot shows unregulated prediction from the ecModel that works on optimality principles and the regulation predictions from the hybrid model. (B) The regulated model forces the use of isoenzymes. Here we display the regulated (Hybrid model), the unregulated (ecModel) and the proteomics data for the isoenzymes catalysing the HXK reaction. In the lower panel the the core reactions in the metabolism under (C) respiration and (D) fermentation are shown. The fluxes are represented by the width of the connectors where dotted lines represent zero flux. The colour of the connectors represent the change in flux from the unregulated ecModel compared to the regulated hybrid model. The FCCs are represented in the figure where the unregulated case is depicted by circles and compared to the regulated case depicted in squares.

Pathway enrichment analysis of the proteins miss-predicted by more than one order of magnitude, by the hybrid model has been performed and showed that the superpathway of glucose fermentation was underpredicted for both respiration and fermentation (p-value of 1.398426e-7 and 7.002912e-5, respectively). Additionally, the superpathway of TCA cycle and glyoxylate cycle was underpredicted (p-value = 0.036267), whilst aerobic respiration and electron transport chain were significantly overpredicted (p-value = 2.857451e-23) in the fermentative state and the pentose phosphate pathway (p-value= 2.863892e-4) as well as glucose-6-phosphate biosynthesis (p-value= 0.019529) were underpredicted in respiration. A detailed comparison between the models as well as in-depth results from the protein predictions are available in Data file S1 and Supplementary Information S1.

To better understand which pathways and reactions are most affected by regulation, the metabolic flux distributions predicted by the hybrid model and the ecModel were compared. Larger flux differences arose for respiratory conditions, in which the average relative change in flux was 1.85 in contrast to 0.46 in fermentation (Data file S2), this result is heavily influenced by the amount of totally activated or deactivated fluxes by the hybrid model, 57 for respiration and 29 for fermentation (Figure 3, Data file S2). ecModels provide a network in which reversible metabolic reactions are split, creating separate reactions for the forward and backwards fluxes, thus distributions of net metabolic fluxes were also obtained and compared among models and conditions. As some enzymes are upregulated by the hybrid model even to levels that exceed the flux capacity of certain pathways (for a fixed growth rate), futile fluxes are expected to arise across the metabolic network.

Increased exchange fluxes for glucose, oxygen and acetate were observed in the respiration phase (Data file S2). Additionally, an increase in the overall flux through the pentose phosphate pathway as well as an induced use of NDE instead of only NDI, allowing for the utilization of cytosolic NADH to reduce oxygen demands in the oxidative phosphorylation pathway was detected and has previously been associated with chronological ageing (Wei Li et al., 2006). Increased flux on PCK, PDC, ALD2 and ACS, around the pyruvate branching point, led to an overall increased flux through the TCA cycle (Figure 3 and Data file S2). To balance the increased production of AMP by ACS the ADK reaction is also upregulated. Futile fluxes are induced by regulation in galactose metabolism (GAL7 and GAL10), lower glycolysis (TDH, PGK and ENO), TCA cycle (FUM and MDH), as well as TKLa and PGM in the pentose phosphate pathway and ADH (Figure 3 and Data file S2) in respiration.

In the fermentation state, futile fluxes also occur in galactose metabolism (GAL7) as well as glycolysis (PGI, PGK, TPI and GPM), TAL1 in the pentose phosphate pathway and ACO in the TCA cycle (Figure S4 and Data file S2). Down-regulation of oxidative metabolism, increased uptake of glucose and increased flux through glycolysis were observed, which is consistent with the changes that have been attributed to glucose-induced repression during the long term Crabtree effect (de Alteriis et al., 2018) (Figure 3).

The control exerted by each enzyme on the global glucose uptake rate was investigated through the calculation of flux control coefficients (FCCs), allowing comparison of the distribution of metabolic control between the pure enzyme-constrained and hybrid model. In both conditions the FCCs obtained for hexokinases by the ecModel (HXK1 for respiration and HXK3 in fermentation) showed a value equal to 1, the highest value in their respective distributions, indicating that the overall glucose uptake rate is mostly governed by the activity of this enzymatic reaction step. In contrast, the constraints applied by the hybrid model distribute the control over the glucose uptake flux in a more even way across different enzymes and pathways, yielding FCCs of 0 for the different HXK isoforms in both metabolic regimes.

As a general trend, more FCCs with a high value (FCC>0.05) are obtained for fermentative conditions than for respiration, despite the use of the ecModel or hybrid model (Figure 3 and Data file S3). In the respiratory condition, the highest FCCs are concentrated in the oxidative phosphorylation pathway as well as around the branching point of pyruvate and PFK in glycolysis, whose activity is related to the connections between glycolysis and PP pathway. Moreover, the absence of glucose uptake control by lower glycolytic enzymes, and the prevalence of a non-zero FCC for PFK in both ecModel and hybrid model agrees with experimental evidence for mouse cell-lines in respiratory conditions (Tanner et al., 2018). For the fermentative condition the highest FCCs are concentrated in the TCA cycle. Similarly to the respiration case, non-zero FCCs are present in the reaction steps surrounding the connecting points of different pathways, such as PFK, FBA and TDH connecting glycolysis with the pentose phosphate pathway and reactions around pyruvate, which connect glycolysis with fermentation and the TCA cycle (Figure 3), this trend might indicate that in these branching points kinetic control is still a relevant mechanism governing fluxes.

### Deletion of the Snf1 in the hybrid model shows the importance of the Snf1 pathway in low glucose conditions and attributes the connection between regulation and chronological ageing to Snf1

To investigate how the individual signaling pathways contribute to changes in metabolic fluxes, the main component of each signaling pathway was deleted and flux changes were compared between the wild-type hybrid model and the knockout versions (Data file S4).

The Snf1 deletion was the only deletion showing any effect on the net fluxes in the respiratory condition (Figure S4) while the Reg1, PKA and TOR deletions showed effects in fermentation conditions, consistent with the deletion experiments done with the Boolean model. The different mutants in fermentation do not induce major changes in net fluxes, however, the enzyme usage profile differs across the different mutants. Notably, the largest changes in terms of futile fluxes were observed in the TPI reaction, repressed in respiration by the Snf1 pathway and activated in fermentative conditions by either the PKA or Reg1 pathways. In respiration, Snf1 is also responsible for the futile fluxes through GPM, PGI and reduces the futile fluxes trough FUM, MDH, PGM and GAL10. The model simulations show a less diverse use of isoenzymes in all knockouts, which is most likely due to the reduction in the complexity of the regulatory layer. Considering the inherent property of flux balance analysis, any reduction in the regulatory network will be closer to the optimal distribution in which just the most efficient isoforms are used.

The Snf1 deletion exhibits an overall decrease in the flux towards respiration and a large decrease in flux through PPP, showing also a relatively strong downregulation of enzymatic steps surrounding pyruvate. The most significant changes are observed in NDE and PCK that are turned off and ALD6 which is turned on, implying that the Snf1 pathway is responsible for changing the acetate production via ALD6 to acetate production via ALD2, resulting in increased production of cytosolic NADH to the expense of the NADPH, which is compensated by increasing the flux through the pentose phosphate pathway as well as the additional use of NDE.

## Discussion

The effects of nutrient-induced signaling on metabolism play an important role in maintaining organismal homeostasis and consequently understanding human disease and ageing. To gain a better understanding of the interconnectivity between nutrient signaling and metabolism, we have developed a hybrid model by combining the Boolean and the enzyme constraint models using a regulatory network as a link. More specifically, we have implemented a Boolean signaling network that is responsive to glucose and nitrogen and an ecModel of yeast’s central carbon metabolism. The proposed framework has been validated using available experimental data resulting in an increased predictive power on individual protein abundances in comparison to individual models alone. Further we were able to characterize the cells deviation from the optimal protein allocation and flux distribution profiles. The model is capable of reproducing the regulatory effects that are associated with the Crabtree effect and glucose repression. The model showed a connection between SNF1 signaling and chronological lifespan by the regulation of NDE and NDI usage in respiring conditions. In addition, the model showed that during fermentation, enzyme utilization is the more important factor governing the protein allocation, while in low glucose conditions robustness and control is prioritized.

The integration of regulation constraints is resulting in a highly constrained hybrid model. The downside of this approach is connected to the lack of information regarding the regulatory effects of transcription factor activation. In this work we assume a uniform proportional action for all gene targets, together with the other constraints of the model, resulting in a rather low effect on the regulatory action. Despite this, the hybrid model shows improved protein abundance predictive power and can qualitatively reproduce regulatory effects associated with glucose repression in fermentation conditions, suggesting that with this framework we can gain novel insight into the interplay between signaling pathways and metabolism, however, any quantitively or definitive statements should be avoided. Another limitation is the inclusion of only the central carbon metabolism, a potential extension of this work would include the addition of pathways responsive to glucose signaling, like glycerol metabolism and fatty acid synthesis, enabling also the study of the regulatory effect on these pathways specifically with relatively few modifications in the hybrid model.

The hybrid model shows that under regulation the NADH to support the electron transport chain is partly coming from the cytosol with the help of the mitochondrial external NADH dehydrogenase, NDE2. Overexpression of NDI1, in contrast to NDE1, causes apoptosis-like cell death which can be repressed by growth on glucose-limited media (Wei Li et al., 2006). In our model regulation acts on both NDE and NDI which will lower the need for NDI1 expression and thus causing apoptosis-like cell death. The hybrid model gives the ability to determine that the Snf1 pathway alone is responsible for the shift to the additional use of NDE and NDI instead of only NDI. Snf1 is active in glucose-limited media and thus would help mitigate the phenotype of over expressed NDI1. With these result, we can connect Snf1 with the respiration-restricted apoptotic activity described by Wei Li et al and help explain how Snf1 is connected to chronological ageing (Wierman et al., 2017).

Futile fluxes in the cell have been examined previously within the constraints of osmotics, thermodynamics and enzyme utilization (Park et al., 2016b), where the osmotics are putting a ceiling on the allowed metabolite concentrations in the cell while thermodynamics govern the net fluxes through reactions. The induced futile fluxes can be explained by the fact that regulation included in the hybrid model will force the cell to use some enzymes even above its pathway flux requirements, adding robustness of metabolism to a constantly changing environment. The increase in flux in both forward and backward directions (i.e the increased futile flux through reactions) implies that these enzymes are working closer to their equilibrium and thus have a low flux control over the pathway flux, while enzymes with a strong forward flux have large flux control (Kacser et al., 1995). This feature is also displayed by our hybrid model, in which all enzymatic steps with induced futile fluxes exert null control over glucose uptake (FCCs = 0). More enzymes in a pathway working close to their equilibrium results in robustness against perturbations as well as allows the pathway to be controlled and regulated through a few enzymes, however, this happens at the expense of inefficient utilization of enzymes as the cell needs to spend more resources to sustain a pool of enzymes that are carrying both forward and backward fluxes (Noor et al., 2016; Park et al., 2016a). Our predictions of several glycolytic steps forced to operate closer to their equilibrium by regulation (high futile fluxes induced for TDH, PGK and ENO in respiration, and for TPI, PGK and GPM in fermentation) agree with experimental studies on *E. coli*, iBMK cells and *Clostridia cellulyticum*, which have suggested the utility of near-equilibrium glycolytic steps not just for providing robustness to environmental changes but also for enhancing metabolic energy yield (Park et al., 2019).

Computation of FCCs showed that in respiration the glucose flux is tightly dependent on the activity of the enzymatic steps in oxidative phosphorylation, a high-energy yield pathway. In contrast, in the fermentative condition flux control is split between PFK, PYK, PDC and several steps in the TCA cycle. Interestingly, the FCCs in TCA cycle are decreased by around the half after applying the regulatory constraints in the hybrid model, providing hints of the importance of enhancing robustness in this pathway at high growth rates due to increased demand of biomass precursors. The prevalence of the highest FCCs in fermentation for PFK, PYK and PDC (for both ecModel and hybrid model) indicate their important role as modulators of flux balance between glycolysis, PPP and fermentative pathways at highly demanding conditions, suggesting that when entering fermentation, the cell sacrifices robustness to favor enzyme utilization.

Comparison of enzyme usage and flux distributions between models and across conditions reveals that the effects of regulation are generally stronger for the respiratory condition, causing the arisen of more and higher futile fluxes; turning on reaction steps that are not required by optimal metabolic allocation (purely ecModel); and inducing higher fold-changes into fluxes. These findings suggest that metabolic phenotypes are majorly shaped by regulatory constraints in low glucose conditions, whilst enzymatic constraints play a major role when glucose is not the limiting resource.

It was also found that the regulatory layer diminishes the strong flux control that hexokinase isoforms have over glucose consumption in both low and high glucose conditions to 0. The hexokinases in yeast, specially HXK2, have a central role in glucose signaling. It works both as an effector in the Snf1 pathway and also actively participates in the repression complex together with Mig1 in glucose repression during high glucose conditions (Vega et al., 2016). Intuitively, it would be practical if an enzyme having these central and diverse tasks in the cell would not have such a high FCC as can be seen with the ecModel. When small perturbations in enzyme activity or concentration have large effects on glucose consumptions, allocating this enzyme to other parts of the cell such as the nucleus while participating in the repression complex, would be energetically expensive. Given the central role of hexokinase in glucose signaling, this would be of interest for further investigation and future studies.

Overall, in this work, we have shown how the hybrid modeling framework integrating nutrient-sensing pathways and central carbon metabolism can not only improve individual model predictions but can also elucidate how single components in the dynamic signaling layer affect the steady-state metabolism. We tested our model against both respiring and fermenting conditions and could not only predict known phenomena but also find novel connections. This methodology can be used to connect both original and readily available models in yeast to look at the interactions between signaling and metabolism. This can be applied to genome-scale and on different subsystems of metabolism and for different signaling systems, for example, macronutrients or osmotic stress etc. The availability of genome-scale models of different organisms is constantly growing and with our increasing understanding of signaling systems and regulatory networks, the methodology developed in the course of this work can be adapted to many other organisms. Hybrid models, like the one proposed here, provides a framework for testing a different hypothesis, as we demonstrated by knocking out several components of the nutrient-induced signaling network. In summary, we developed a methodology to investigate intrinsically different systems, such as signaling and metabolism, in the same model gaining insight into how the interplay between them can have non-trivial effects.

## Materials and methods

### 1.1 Boolean model of nutrient-induced signaling pathways

Based on an extensive literature review, a detailed topology of the nutrient-induced signaling pathways TORC1, SNF1 and PKA accounting also for their crosstalks was derived and formalized as a Boolean network model using a vector-based modelling approach (Welkenhuysen et al., 2019) **TORC1**: (Broach, 2012; Fernández-García et al., 2012; Hong et al., 2003; Kacherovsky et al., 2008; Leverentz & Reece, 2006; Ludin et al., 1998; MacPherson et al., 2006; Santangelo, 2006; Sanz et al., 2000; Schüller, 2003; Smith et al., 2011; Soontorngun et al., 2012; Sutherland et al., 2003; Turcotte et al., 2010; Westholm et al., 2008; Woods et al., 1994); **SNF1:** (Broek et al., 1987; Colombo et al., 1998; Dihazi et al., 2003; Hu et al., 2010; Jones et al., 1991; Kataoka et al., 1985; Kraakman et al., 1999; Ma et al., 1999; Martínez-Pastor et al., 1996; Matsumoto et al., 1982; Nikawa et al., 1987; Pedruzzi et al., 2000; K. Peeters et al., 2017; T. Peeters et al., 2006; Portela et al., 2002; Rittenhouse et al., 1987; Robinson et al., 1987; Rolland et al., 2000; Sass et al., 1986; Schepers et al., 2012; Swinnen et al., 2006; K Tanaka et al., 1989, 1990; Kazuma Tanaka et al., 1990; T Toda et al., 1987; Takashi Toda et al., 1985, 1987); **PKA:** Bar-Peled et al., 2013; Beck & Hall, 1999; Binda et al., 2009; Bonfils et al., 2012; Broach, 2012; Conrad et al., 2014; Dilova, Aronova, Chen, & Powers, 2004; Dubouloz, Deloche, Wanke, Cameroni, & De Virgilio, 2005; Georis, Feller, Vierendeels, & Dubois, 2009; Hughes Hallett, Luo, & Capaldi, 2014; Kuruvilla, Shamji, & Schreiber, 2001; Lempiäinen et al., 2009; Liu & Butow, 1999; Marion et al., 2004; Reinke et al., 2004; Urban et al., 2007; Wanke et al., 2008; Yan, Shen, & Jiang, 2006; **crosstalks**: Barrett, Orlova, Maziarz, & Kuchin, 2012; Castermans et al., 2012; Cherry, Johnson, Dollard, Shuster, & Denis, 1989; Hughes Hallett et al., 2014; Nicastro et al., 2015; Wanke et al., 2008).

The model consists of four different components: metabolites, target genes, regulated enzymes and proteins. For the regulated enzymes, presence and phosphorylation state were considered whereas metabolites and target genes were only described by a single binary value indicating their presence and transcriptional state respectively. The state vectors were translated into a single binary value indicating the components’ activity, allowing a better graphical depiction. In total, the model comprises 5 metabolites, 10 groups of target genes, 6 enzymes whose activity is altered upon nutrient signaling and 46 proteins belonging to PKA/cAMP, the SNF1 and the TORC1 pathway, for detailed description, see Supplementary Information S1 and Table S1-S6.

Availability of glucose and nitrogen was used as an input to the model and is implemented as one vector of binary values for each nutrient. This input enables to simulate the induction of signaling under different nutrient conditions, for instance the addition of glucose and nitrogen to starved cells is represented by the vector 0|1 for both nutrients respectively. Here, 0 represent the starved or low nutrient condition and 1 the nutrient-rich condition. Based on this input and the formulation of the Boolean rules, a cascade of state transitions is induced. The simulation was conducted using a synchronous updating scheme meaning that at each iteration, the state vectors are updated simultaneously. The algorithm stops if a Boolean steady state is reached at which no operation causes a change in the state vectors. This process is repeated for each pair of glucose and nitrogen availabilities whereby the reached steady state for each nutrient condition serves as an initial condition for the next nutrient condition.

Since for many of the included processes, no information on the mechanisms causing reversibility was available, especially a lack in the knowledge on phosphatases reverting phosphorylation was observed (Welkenhuysen *et al* 2019), gap-filling was conducted by including else-statements. This ensures that a component’s state vector changes again e.g. if the conditions causing its phosphorylation are not fulfilled anymore. This gap-filling process guarantees the functionality of the Boolean model in both directions, meaning the simulation of state transitions occurring when nutrients (glucose and nitrogen) are added to nutrient-depleted cells as well as when cells are starved for the respective nutrients. Crosstalk mechanisms between the pathways were formulated as if-statements and can be switched off (0) or on (1). Furthermore, a simulation of knockouts of the pathways’ components is possible by setting the value indicating their presence to 0.

#### Enzyme-constrained metabolic model

A reduced stoichiometric model of *Saccharomyces cerevisiae’s* central carbon and energy metabolism, including metabolites, reactions, genes and their interactions accounting for glycolysis, TCA cycle, oxidative phosphorylation, pentose phosphate, Leloir and anaerobic excretion pathways, together with a representation of biomass formation, was taken as a network scaffold(Nilsson & Nielsen, 2016). The metabolic model was further enhanced with enzyme constraints using the GECKO toolbox v1.3.5 (Sánchez et al., 2017), which considers enzymes as part of metabolic reactions, as they are occupied by metabolites for a given amount of time that is inversely proportional to the enzyme’s turnover number (*k*_*cat*_). Therefore, enzymes are incorporated as new “pseudo metabolites” and usage pseudo reactions are also introduced in order to represent their connection to a limited pool of protein mass available for metabolic enzymes. Moreover, all reversible reactions are split into two reactions with opposite directionalities in the ecModel, in order to account for the enzyme demands of backwards fluxes. Several size metrics for the Boolean model, the metabolic network and its enzyme-constrained version (ecModel) are shown in Table 1.

**Table 1.** Size metrics for the Boolean, original metabolic model and its enzyme-constrained version.

| Boolean model |  |
| --- | --- |
| Metabolites | 5 |
| Target gene groups | 10 |
| Enzyme PTMs | 6 |
| Proteins | 46 |
| Metabolic model |  |
| Reactions | 90 |
| Metabolites | 81 |
| Genes | 130 |
| Cellular compartments | 4 |
| ecModel |  |
| Reactions | 324 |
| Metabolites | 111 |
| Enzymes | 127 |
| Promiscuous enzymes | 41 |
| Reactions with isoenzymes | 30 |
| Enzyme complexes | 11 |
| Reactions w/Kcat | 115 |

As the obtained ecModel has the same structure as any metabolic stoichiometric model, in which metabolites and reactions are connected by a stoichiometric matrix, the technique of flux balance analysis (FBA) can be used for quantitative prediction of intracellular reaction fluxes (Orth et al., 2010). FBA assumes that the metabolic network operates on steady-state, i.e. no net accumulation of internal metabolites, due to the high turnover rate of metabolites when compared to cellular growth or environmental dynamics (Varma & Palsson, 1994), therefore, by setting mass balances around each intracellular metabolite a homogenous system of linear equations is obtained. The second major assumption of FBA is that metabolic phenotypes are defined by underlying organizational principles, therefore an objective function is set as a linear combination of reaction fluxes which allows to obtain a flux distribution by solving the following linear programming problem

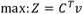

Subject to

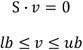

Where *C*^*T*^, is a transposed vector of integer coefficients for each flux in the objective function (*z*); *v*, is the vector of reaction fluxes; *S*, is a stoichiometric matrix, representing metabolites as rows and reactions as columns; *lb* and *ub* are vectors of lower and upper bounds, respectively, for the reaction fluxes in the system. Additionally, the incorporation of enzyme constraints enables the connection between reaction fluxes and enzyme demands, which are constrained by the aforementioned pool of metabolic enzymes

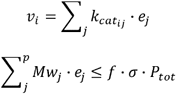

Where 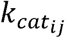 is the turnover number of the enzyme *j* for the i-th reaction, as in some cases several enzymes can catalize the same reaction (isoenzymes); *e*_*j*_, is the usage rate for the enzyme *j* in mmol/gDw h^-1^; *Mw*_*j*_, represents the molecular weight of the enzyme *j*, in mmol/g; *P*_*tot*_, is the total protein content in a yeast cell, corresponding to a value of 0.46 g_prot_/gDw (Famili et al., 2003); *f*, is the fraction of the total cell proteome that is accounted for in our ecModel, 0.1732 when using the integrated dataset for *S. cerevisiae* in paxDB as a reference (M. Wang et al., 2015b); and *σ* being an average saturation factor for all enzymes in the model.

This simple modelling formalism enables the incorporation of complex enzyme-reaction relations into the ecModel due to its matrix formulation, such as isoenzymes, which are different enzymes able to catalyse the same reaction; promiscuous enzymes, enzymes that can catalyze more than one reaction; and enzyme complexes, several enzyme subunits all needed to catalyse a given reaction.

#### ecModel curation

As the ecModel was generated by the automated pipeline of the GECKO toolbox, several of its components were curated in order to achieve predictions that are in agreement with experimental data at different dilution rates. Data on exchange reaction fluxes at increasing dilution rates, spanning both respiration and fermentative metabolic regimes (Van Hoek et al., 1998) was used as a comparison basis. Additionally, all unused genes in the original metabolic network were removed and gene rules for lactose and galactose metabolism were corrected according to manually curated entries for *S. cerevisiae* available at the Swiss-Prot database (Bateman, 2019). Gene rules and metabolites stoichiometries (P/O ratio) in the oxidative phosphorylation pathway were also corrected according to the consensus genome-scale network reconstruction, Yeast8 (Lu et al., 2019).

The average saturation factor for the enzymes in the model was fitted to a value of 0.48, which allows the prediction of the experimental critical dilution rate (i.e. the onset of fermentative metabolism) at 0.285 h^-1^. ATP requirements for biomass production were fitted by minimization of the median relative error in the prediction of exchange fluxes for glucose, oxygen, CO_2_ and ethanol across dilution rates (0 – 0.4 h^-1^), resulting in a linear relation depending on biomass formation from 18 to 25 mmol per gDw for respiratory conditions and from 25 to 30 mmol per gDw for the fermentative regime.

#### Hybrid model

A hybrid model consists of the Boolean model connected with the ecModel trough a transcriptional layer that regulates its constraints on protein allocation (Figure 1). The active transcription factors act on the upper or lower bounds of the enzyme usage pseudo reaction depending on down- or up regulation, respectively. The magnitude of the induced perturbations is calculated according to previously calculated enzyme usage variability ranges, subject to a given growth rate and optimal glucose rate, expressed as

Upregulation:

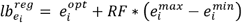

Downregulation:

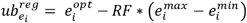

Where 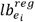 and 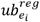 represent the lower and upper bounds for the usage pseudoreaction of enzyme *i* in the regulated model; 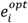, is a parsimonious usage for enzyme *i* for a given growth and glucose uptake rates; *RF*, corresponds to a regulation factor between 0 and 1; 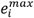 and 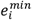 are the maximum and minimum allowable usages for enzyme *i* under the specified conditions.

A distribution of parsimonious enzyme usages is obtained by applying the rationale of the parsimonious FBA technique (Lewis et al., 2010), which explicitly minimizes the total protein burden that sustains a given metabolic state (i.e. fixed growth and nutrient uptake rates).

To connect the transcription factor activity with gene regulation we extracted regulation information from YEASTRACT and set a regulation level of 5% of the enzyme usage variability range for the simulations. When several transcription factors affect the same gene, the effects are summed up and the resulting sum is used as a basis for constraint. For example, if a gene is downregulated by two transcription factors (−2) and upregulated by one transcription factor (+1), the net sum would be (−1), thus the gene will be downregulated. In our model, an absolute sum higher than 1 will not cause a stronger regulation, as this additive process is just implemented to define the directionality of a regulatory effect.

### 2.5 Enzyme usage variability analysis

As metabolic networks are highly redundant and interconnected, the use of purely stoichiometric constraints usually leads to an underdetermined system with infinite solutions (Kauffman et al., 2003), in a typical FBA problem it is common that even for an optimal value of the objective function, several reactions in the network can take any value within a “feasible” range, such ranges can be explored by flux variability analysis (Mahadevan & Schilling, 2003).

In this study, enzyme usage variability ranges for all of the individual enzymes are calculated by fixing a minimal glucose uptake flux, for a given fixed dilution rate, and then running sequential maximization and minimization for each enzyme usage pseudo reaction.

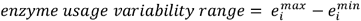

Subject to:

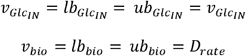

This approach allows the identification of enzymes that are either tightly constrained, highly variable or even not usable at optimal levels of biomass yield.

#### Simulations

Cellular growth on chemostat conditions, at varying dilution rates from 0 to 0.4 h^-1^, was simulated with the multiscale model by the following sequence of steps:

1. Initially, the desired dilution rate is set as both lower and upper bounds for the growth pseudo reaction and the glucose uptake rate is minimized, assuming that cells maximize biomass production yield when glucose is limited (Oliveira et al., 2005; Schuetz et al., 2007)

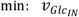

Subject to

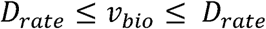
2. The obtained optimal uptake rate 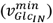 is then used as a basis to estimate a range of uptake flux to further constrain the ecModel.

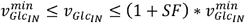

As 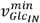 represents the minimum uptake rate allowed by the stoichiometric and enzymatic constraints of the metabolic network, possible deviations from optimal behaviour may be induced by regulatory circuits. In order to allow the Boolean model to reallocate enzyme levels a suboptimality factor (sF) of 15% was used to set an upper bound for 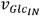.
3. The ecModel is connected to the glucose-sensing Boolean model through the glucose uptake rate. At the critical dilution rate, the glucose uptake rate obtained by the ecModel is 3.2914 mmol/gDw h, this value is used as a threshold to define a “low” or “high” glucose level input in the Boolean model, represented as 0 and 1, respectively. For each dilution rate, the initial value of 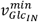 is calculated and fed to the regulatory network, which runs a series of synchronous update steps until a steady-state is reached.
4. At steady state, the regulatory network indicates the enzyme usages that should be up and downregulated, for which new usage bounds are set as described above.
5. A final FBA simulation is run by minimizing the glucose uptake rate, subject to a fixed dilution rate and the newly regulated enzyme usage bounds.

Gene deletions can also be set in the Boolean module and will result in activation or inactivation of transcription factors which then affect the constraints on the FBA model. We ran four simulations of deletion strains as follows: TOR1 and TOR2 (TOR deletion), Snf1 (SNF1 deletion), Tpk1, Tpk2 and Tpk3 (PKA deletion) and Reg1(Reg1 deletion).

#### Proteomics analysis

Protein abundance data on respiratory and fermentative conditions were compared to protein usage predictions by the hybrid model in order to assess its performance. For the respiration phase, absolute protein abundances were taken from a study of yeast growing under glucose-limited chemostat conditions at 30°C on minimal mineral medium with a dilution rate of 0.1 h^−1^ (Doughty et al., 2020).

For the fermentation phase, a proteomics dataset was taken from a batch culture using minimal media with 2% glucose and harvested at an optical density (OD) of 0.6 (Paulo et al., 2016). The dataset given as relative abundances was then rescaled to relative protein abundances in the whole-cell according to integrated data available for *S. cerevisiae* in PaxDB (M. Wang, Herrmann, Simonovic, Szklarczyk, & von Mering, 2015), and finally converted to absolute units of mmol/gDw using the “total protein approach” (Wiśniewski & Rakus, 2014).

We used three metrics for comparing the simulations with the proteomics data, the Pearson correlation coefficient (PCC), two-sample Kolmogorov-Smirnov (KS) test and the mean of the absolute log_10_-transformed ratios between predicted and measured values (r). The PCC and the significance of the PCC were determined by a permutation test of n=2000. Pathway enrichments were done using YeastMine (Balakrishnan et al., 2012) with the Holm-Bonferroni test correction and a max p-value of 0.05.

#### Flux control coefficients

In order to investigate the relationship between enzyme activities and a given metabolic flux, control coefficients can be calculated for each enzyme in the model according to the definition given by metabolic control analysis (MCA) (Kacser et al., 1995):

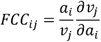

In which 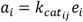 represents the activity of the *i-th* enzyme and *v*_*j*_ is the flux carried by the *j-th* reaction. These coefficients represent the sensitivity of a given metabolic flux to perturbations on enzyme activities, providing a quantitative measure on the control that each enzyme exerts on specific fluxes.

As ecModels include enzyme activities explicitly in their structure, flux control coefficients can be approximated by inducing small perturbations on individual enzyme usages:

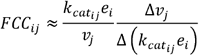

In our hybrid model, perturbations on individual enzyme usages (*e*_*i*_) are induced in relation to a parsimonious usage 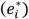 which is compatible with a given set of constraints

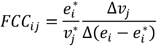

Perturbations equivalent to 0.1% of the parsimonious usage are used for each enzyme. For those cases in which the previously applied constraints do not allow such modification in a given enzyme usage, their activity is then perturbed by operating on the corresponding turnover number for the enzyme-reaction pair 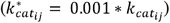 in order to simulate a perturbation in their overall activity.

The model, code and datasets used for this study can be found in the GitHub repository YeastHybridModelingFramework https://github.com/cvijoviclab/YeastHybridModelingFramework.

## Acknowledgments

We would like to thank members of the Hohmann, Nielsen and Cvijovic labs for valuable input. Special thanks to Avlant Nilsson for his contributions to the curation of the original metabolic network used in this study and valuable discussions on the role of enzyme constraints.

## Supplorting Information

SI. Supporting information. Includes detailed description of mechanisms reflected in the Boolean model of nutrient signalling.

