## Supplementary file 1 for "A novel yeast hybrid modeling framework integrating Boolean and enzyme-constrained networks enables exploration of the interplay between signaling and metabolism"

### 1 Detailed description of mechanisms reflected in the Boolean model of nutrient signaling

#### 1.1 SNF1

Snf1 activation upon glucose depletion is associated with increased phosphorylation of Thr210 mediated by the upstream kinases Tos3, Sak1 and Elm1 which are partially redundant, this activation appears to be constitutively (Hong et al. 2003; Sutherland et al. 2003). The return to the inactive state has been attributed to the Reg1-Glc7 complex where studies show that Reg1 interacts with Snf1 (Ludin, Jiang, and Carlson 1998) and targets Glc7 to the complex(Sanz et al. 2000). The model suggest that Reg1-Glc7 binds to Snf1p, mainly relevant in low glucose conditions, and Snf1p phosphorylates Reg1, Glc7 dephosphorylates Reg1. Hxk2 is then either 1) promoting binding of Reg1 to Snf1, 2) promoting phosphorylation of Reg1 or 3) interfering with dephosphorylation by Glc7. In response to high glucose Reg1-Glc7 is dephosphorylating and thus releasing Snf1 from the complex. The dephosphorylation of Reg1 appears to increase efficiency of Glc7 dephosphorylation as well as allowing Reg1 to be released from SNF1(Sanz et al. 2000). This complex form of regulation is a highly adaptable system with a fast response. In our model, this is implemented in a way that when glucose is absent Snf1 is phosphorylated by the upstream kinases. The SNF1 complex phosphorylates Reg1-Glc7. To account for the involvement of Hxk2 in our model we chose to implement mechanism number 2) where Hxk2p can phosphorylate Reg1 (Fernández-García et al. 2012). Phosphorylated Reg1 obstructs the activation of Snf1 and activates Glc7. In this low glucose state SNF1 is active, Reg1 and Hxk2 are phosphorylated and Glc7 is active. When glucose is added, Hxk2 is unphosphorylated and a “high glucose signal”, in our model mediated trough the PKA pathway (Barrett et al. 2012; Castermans et al. 2012) allows Glc7 to dephosphorylate Snf1. When Snf1 and Hxk2 are no longer phosphorylated Reg1 gets dephosphorylated by Glc7 and also Glc7 becomes inactive. In this high glucose condition Snf1 and Reg1 are unphosphorylated and Glc7 is inactive. SNF1-mediates phosphorylation of the transcriptional factors Mig1, Cat8, Sip4 and Adr1 as well as directly phosphorylates and inactivates ACC1(Woods et al. 1994). Mig1 is a repressor which is active in high glucose conditions and represses genes used for alternative carbon sources, mainly SUC, MAL and GAL genes (Westholm et al. 2008; Santangelo 2006; Broach 2012; Schüller 2003). In absence of glucose, Cat8 and Sip4 are activating the transcription of genes regulated by carbon source-responsive elements (CSRE) such as FBP1, PCK1 and ICL1 (Broach 2012; Leverentz and Reece 2006; Turcotte et al. 2010; MacPherson, Larochelle, and Turcotte 2006). Adr1 induces genes involved in use of alternative carbon sources such as ADH1, ACS1 and GUT1 as well as peroxisome biogenesis and fatty acid utilization such as POX1 and PXA1(Turcotte et al. 2010; Soontorngun et al. 2012; Broach 2012; Kacherovsky et al. 2008; Smith et al. 2011) It has been shown that PKA can inactivate the Adr1(Cherry et al. 1989).

#### 1.2 PKA pathway

The protein kinase A (PKA)/cAMP pathway mainly represses genes involved in stress tolerance and post diauxic growth when glucose is available. This means that properties associated with slow, resperative growth and stationary phase are negatively regulated by glucose (Conrad et al. 2014). Intra- and extracellular glucose sensing is carried out by two distinct G-protein systems, namely the Ras pathway and the Gpr1/Gpa2 pathway (Rolland et al. 2000). Ras proteins are small monomeric GTP-binding proteins that are regulated through a cycle of GDP/GTP exchange and GTP hydrolysis. This process is regulated by Cdc25 that triggers the exchange from GDP to GTP on the one hand (Jones, Vignais, and Broach 1991; Robinson et al. 1987; Broek et al. 1987) and by Ira1 and Ira2 that stimulate GTP hydrolysis on the other hand(K Tanaka et al. 1990; Kazuma Tanaka et al. 1990; K Tanaka, Matsumoto, and Toh-E 1989). Sensing of extracellular glucose occurs via the G-protein coupled receptor (GPCR) Gpr1 that interacts with Gpa1. Glucose availability causes a Gpr1-mediated nucleotide exchange in Gpa2 from GDP to GTP yielding its activation (Kraakman et al. 1999; Colombo et al. 1998). Activated Gpa2 as well as/together with activated Ras can stimulate cellular cAMP production via the adenylate cyclase (AC) (Kataoka, Broek, and Wigler 1985; Takashi Toda et al. 1985; Rolland et al. 2000). GPCRs require Ras activation to activate AC which require activity in the upper metabolism such as activity in hexose kinases (Rolland et al. 2000). It has been shown that accumulation of F16BP is coupled to Ras activation (K. Peeters et al. 2017). RAS mutants imitates the AC mutants and the lethality of RAS deletion can be alleviated by *bcy1* mutant cells in the same fashion as AC mutants (Takashi Toda et al. 1985). In the model this is implemented so that Cdc25 is activated by F1,6BP which is present when glucose is present, and Ira is active when no glucose is present. In this model, Ras is able to activate AC but Gpa2 also requires active Ras to activate AC. AC is deactivated by crosstalk with the SNF1 pathway (Nicastro et al. 2015). ´The protein kinase A (PKA) is a heterotetrameric protein complex consisting of two catalytic (Tpk1-3) and two regulatory (Bcy1) subunits(Takashi Toda et al. 1987; T Toda et al. 1987; Matsumoto et al. 1982). Binding of cAMP to the Bcy1 subunits causes the complex to dissociate, thus releasing the blockade of Tpk1-3 kinase activity (Conrad et al. 2014). In contrast, the kelch repeat proteins Krh1 and Krh2 stimulate the association of catalytic and regulatory subunits resulting in an increased amount of cAMP required for PKA activation. However, it was shown that active Gpa2 inhibits Krh activity (T. Peeters et al. 2006). Upon the numerous PKA targets are the phosphodiesterases Pde1 and Pde2 which conciliate a negative feedback mechanism on PKA itself by degrading cAMP (Sass et al. 1986; Nikawa, Sass, and Wigler 1987; Ma et al. 1999; Hu et al. 2010). This is implemented in the model so that the PKA complex is defined as the catalytic subunits and the regulatory subunit is required when the PKA complex is inactive. To activate PKA, the Krh proteins have to be inactive. In the model only phosphorylated Pde breaks down cAMP. PKA also inactivates the Rim15 protein kinase by phosphorylation which is therefore not able to activate the transcription factors Msn2, Msn4 and Gis1 (Swinnen et al. 2006). This can also be achieved through crosstalk with the TOR pathway (Wanke et al. 2008). In an active state, the former two induce expression of genes containing a stress response element (STRE) in their promoter whereas the latter induces transcription of genes comprising a post diauxic shift (PDS) element in their promoter (I Pedruzzi et al. 2000; Martínez-Pastor et al. 1996). PKA also directly phosphorylates cytosolic enzymes such as trehalase (Schepers et al. 2012), phosphofructokinase 2 (Dihazi, Kessler, and Eschrich 2003), pyruvate kinase (Portela et al. 2002) and fructose-1,6-bisphosphatase (Rittenhouse, Moberly, and Marcus 1987).

#### 1.3 TOR pathway

The target of Rapamycin (Tor) kinase complex 1 (TORC1) is not directly involved in glucose sensing; however, glucose availability has been shown to highly influence the activity of TORC downstream targets (Hughes Hallett, Luo, and Capaldi 2014). The strongly conserved TORC1 pathway plays a crucial role in promoting anabolic processes and cell growth in response to nitrogen availability which is probably sensed as the level of intracellular amino acids (Broach 2012). TORC1 comprises either Tor1 or Tor2 kinase in association with Kog1, Lst8 and Tco89 (Reinke et al. 2004) and its activity is regulated by the EGO complex consisting of Ego1, Ego2, Gtr1 and Gtr2 (Dubouloz et al. 2005). Multiple complex nitrogen-sensing mechanisms (Binda et al. 2009; Bonfils et al. 2012; Bar-Peled et al. 2013) lead to physical interaction of EGO with TORC1 resulting in activation of the latter under nitrogen-rich conditions (Binda et al. 2009). Active TORC1 then induces several signaling branches - the Sch9 branch, the Tap42-PPase branch as well as the activation of further transcription factors such as Sfp1 (Urban et al. 2007; Yan, Shen, and Jiang 2006; Marion et al. 2004). TORC1 phosphorylates Sch9(Urban et al. 2007) which then directly phosphorylates Rim15 (Wanke et al. 2008). Tap42 phosphorylation is catalyzed by active TORC1 (Yan, Shen, and Jiang 2006). Phosphorylated Tap42 interacts with TORC1 and associates with the catalytic subunit of type 2A phosphatases (PP2A) like Pph21 or Sit4 and thus inhibits their phosphatase activity (Jiang and Broach 1999; Di Como and Arndt 1996). Dissociation of the complex occurring in case of TORC1 inactivity results in PP2A activation (Beck and Hall 1999). PP2A dephosphorylates its downstream targets such as Gat1, Gln3. (Beck and Hall 1999; Kuruvilla, Shamji, and Schreiber 2001). This regulation is complex and different transcription factors are regulated differently depending on TORC1 stimuli (Georis et al. 2009; Broach 2012; Conrad et al. 2014). In this model we chose a reduced representation where Gat1 and Gln3 are dephosphorylated by active PP2A and either of them induce nitrogen catabolite repression (NCR) genes. Rtg1 and Rtg3 promote the expression of retrograde signaling (RTG) genes whose gene products enable alpha-ketoglutarate production and further processing into glutamine and glutamate to sustain amino acid biosynthesis (Liu and Butow 1999). However, signaling via the RTG branch requires some additional regulation which involves Rtg1, 2 and 3 and the negative regulators Mks1 and Bmh1 and 2. In nitrogen-rich conditions, TORC1 phosphorylates Mks1 that complexes with the Bmh proteins thus sequestering Rtg1 and 3 in the cytoplasm. In contrast, nitrogen depletion causes reduced phosphorylation of Mks1 mediated by PP2A, so that the former complexes with Rtg2, consequently releasing Rtg1 and 3 into the nucleus where they act as transcriptional activators (Dilova et al. 2004; Broach 2012). Again, we chose a reduced representation where TORC1 phosphorylates Mks1 and active PP2A dephosphorylates Mks1. Rtg1,3 is phosphorylated unless dephosphorylated Mks1 and Rtg2 are present. Then Rtg1,3 becomes dephosphorylated in the model and activates RTG transcription. TORC1-mediated phosphorylation of Sfp1 results in the transcription factor's nuclear translocation where it induces ribosomal protein and ribosome biogenesis gene expression (Marion et al. 2004; Lempiäinen et al. 2009). There is also a negative feedback mechanism in which phosphorylated Sfp1 negatively regulates the phosphorylation state of Sch9 (Lempiäinen et al. 2009). However, this negative feedback is not implemented in the model.

The described signaling activity may be valid for conditions in which enough glucose is available, and cells are not exposed to any stress factors. However, under glucose depletion, both branches downstream of TORC1, namely the PP2A and the Sch9 branch, show no or only little activity which is probably caused by crosstalk with the Snf1 pathway (Hughes Hallett, Luo, and Capaldi 2014). In this model it is implemented in that way that phosphorylated Snf1 inhibits TORC1 activity as well as it phosphorylates Tap42.

#### 1.4 Crosstalk

To enable an efficient and fine-tuned adaption to environmental conditions such as different nutrient availabilities, interaction of the induced signaling pathways is required to integrate information. Although we integrated some of these crosstalk mechanisms we considered relevant for the model, many more pathway interactions were reported which highlights the fine-tuned regulation of integrating environmental changes. (Shashkova, Welkenhuysen, and Hohmann 2015)

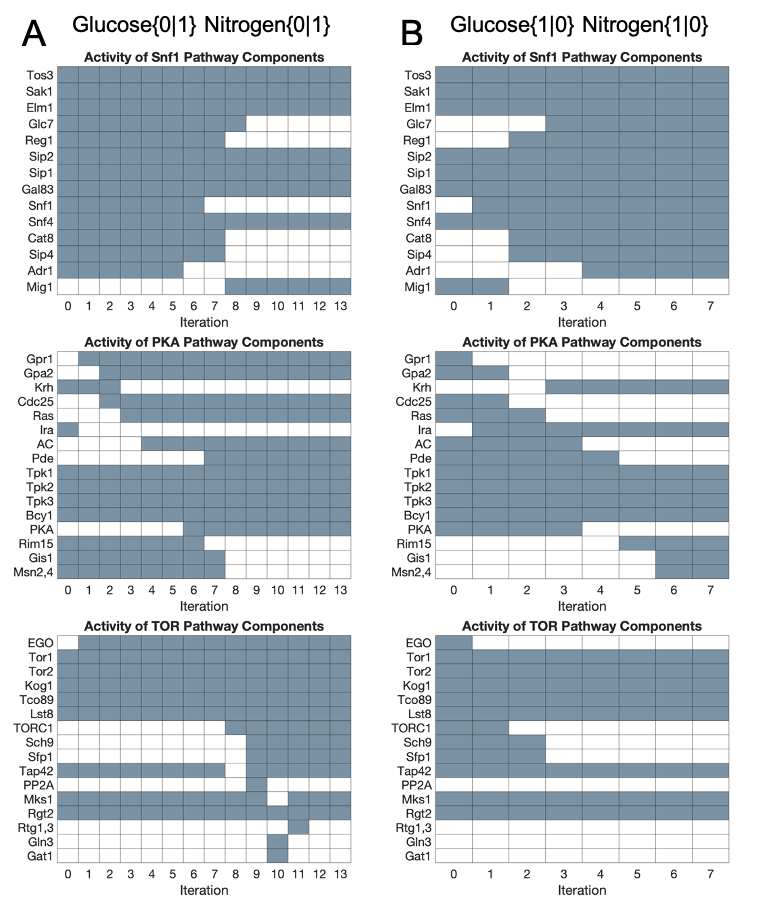

Supplementary Figure 1. Transition map of components in the Boolean model separated by their respective pathway where the blue color indicates activity. Simulations are made with all crosstalk turned on. Panel (A) shows the simulation dynamics going from nutrient depleted conditions to nutrient rich conditions and panel (B) shows the simulation dynamics going from nutrient rich conditions to nutrient depletion.

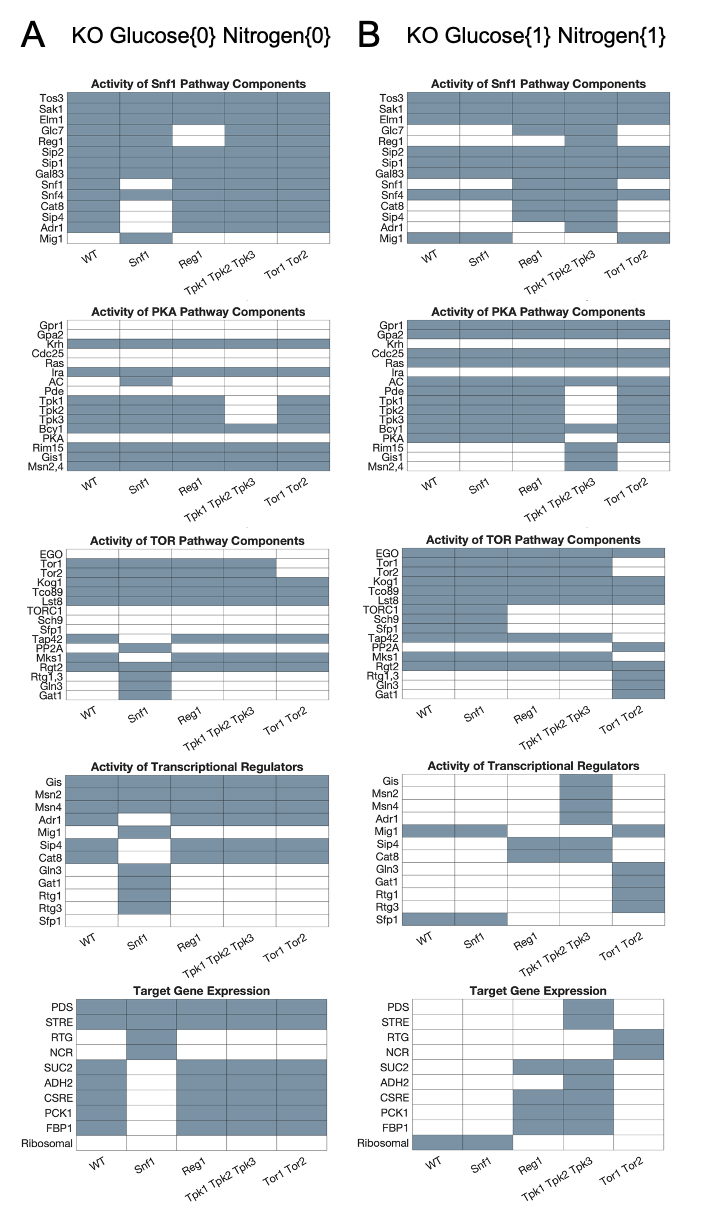

Supplementary Figure 2. Steady state map of components in the Boolean model when knock out (KO) strains are simulated from wild type (WT) to KO where the blue color indicates activity. Panel (A) show the KO behavior in low nutrient conditions compared to the WT and panel (B) show the KO behavior in high nutrient conditions compared to the WT.

Table 1. Rules and references associated to any field of the Boolean vector in the Snf1 pathway.

| SNF1 | | | |
| --- | --- | --- | --- |
| Name | Presence | Phosphorylation | Specific activity |
| Tos3 | Present unless deleted | - | - |
| Sak1 | Present unless deleted | - | - |
| Elm1 | Present unless deleted | - | - |
| Glc7 | Present unless deleted | - | Active when Glc7 is present and Reg1 is present and phosphorylated.  Inactive when Reg1 is unphosphorylated and Glc7 is present or when Glc7 is not present. (Sanz et al. 2000) |
| Reg1 | Present unless deleted | Phosphorylated when Glc7, Reg1, Snf4 and any of Sip2, Sip1 or Gal83 is present as well as Snf1 is phosphorylated and Reg1 is not phosphorylated or if Hxk2 is phosphorylated(Sanz et al. 2000).  Unphosphorylated when Hxk2 and Snf1 is not phosphorylated and Glc7 is present, active and Reg1 is present and phosphorylated(Sanz et al. 2000). | - |
| Sip2 | Present unless deleted | - | - |
| Sip1 | Present unless deleted | - | - |
| Gal83 | Present unless deleted | - | - |
| Snf1 | Present unless deleted | Phosphorylated when glucose is absent, Snf1 is present, any of Tos3, Elm1 or Sak1 is present and, Snf1 and Reg1 is unphosphorylated (Hong et al. 2003; Sutherland et al. 2003; Sanz et al. 2000). Unphosphorylated when Snf1 is not present.  Unphosphorylated when Glucose, Glc7 and Reg1 is present, PKA and Glc7 is active and Reg1 and Snf1 is phosphorylated (Sanz et al. 2000) (Barrett et al. 2012; Castermans et al. 2012) | - |
| Snf4 | Present unless deleted | - | - |
| Cat8 | Present unless deleted | Always unphosphorylated unless Snf1 is active and Snf4 and Cat8 is present as well as any of Sip2, Sip1 or Gal83. (Broach 2012; Leverentz and Reece 2006; Turcotte et al. 2010; MacPherson, Larochelle, and Turcotte 2006). | - |
| Sip4 | Present unless deleted | Always unphosphorylated unless Snf1 is active and Snf4 and Sip4 is present as well as any of Sip2, Sip1 or Gal83. (Broach 2012; Leverentz and Reece 2006; Turcotte et al. 2010; MacPherson, Larochelle, and Turcotte 2006). | - |
| Adr1 | Present unless deleted | Always unphosphorylated unless Snf1 is active and Snf4 and Adr1 is present as well as any of Sip2, Sip1 or Gal83. (Turcotte et al. 2010; Soontorngun et al. 2012; Broach 2012; Kacherovsky et al. 2008; Smith et al. 2011) | - |
| Mig1 | Present unless deleted | Always unphosphorylated unless Snf1 is active and Snf4 and Mig1 is present as well as any of Sip2, Sip1 or Gal83. (Westholm et al. 2008; Santangelo 2006; Broach 2012; Schüller 2003) | - |

Table 2. Rules and references associated to any field of the Boolean vector in the PKA pathway.

| PKA pathway | | | |
| --- | --- | --- | --- |
| Name | Presence | Phosphorylation | Specific activity |
| Gpr1 | Present unless deleted | - | Always inactive unless GLUex and Gpr1 are present. (Kraakman et al. 1999; Colombo et al. 1998). |
| Gpa2 | Present unless deleted | - | Active if Gpr1 is active and Gpa2 is present.  Inactive if Gpr1 is inactive or Gpa2 is deleted.(Kraakman et al. 1999; Colombo et al. 1998). |
| Krh | Present unless deleted | - | Always active unless deleted or Gpa2 is active. (T. Peeters et al. 2006) |
| Cdc25 | Present unless deleted | - | Always inactive unless the metabolite F16BP and pathway component Cdc25 are present. (K. Peeters et al. 2017). |
| Ras | Present unless deleted | - | Active when Ras is present, Cdc25 is active and Ira is inactive (and Ras was previous inactive).  Inactive when Ras is present, Cdc25 is inactive and Ira is active (and Ras is previously active). (Jones, Vignais, and Broach 1991; Robinson et al. 1987; Broek et al. 1987; K Tanaka et al. 1990; Kazuma Tanaka et al. 1990; K Tanaka, Matsumoto, and Toh-E 1989). |
| Ira | Present unless deleted | - | Always inactive unless GLUex and Ira are present. |
| AC | Present unless deleted | - | Active when Ras is active and AC is present, or when Ras and Gpa2 is active and AC is present.(Kataoka, Broek, and Wigler 1985; Takashi Toda et al. 1985; Rolland et al. 2000)  Inactive if AC is deleted.  Inactive when Gpa2 and Ras is inactive, AC, Snf4 is present and Sip2, Sip1 or Gal83 is present and Snf1 is phosphorylated. (Nicastro et al. 2015) |
| Pde | Present unless deleted | Always unphosphorylated unless Pde is present and PKA is active. (Sass et al. 1986; Nikawa, Sass, and Wigler 1987; Ma et al. 1999; Hu et al. 2010). | - |
| Tpk1 | Present unless deleted | - | - |
| Tpk2 | Present unless deleted | - | - |
| Tpk3 | Present unless deleted | - | - |
| Bcy1 | Present unless deleted | - | - |
| PKA | Always absent unless Tpk1, Tpk2 or Tpk3 is present. (Takashi Toda et al. 1987; T Toda et al. 1987; Matsumoto et al. 1982). | - | Active if PKA, Bcy1 and cAMP is present and Krh is inactive or Bcy1 is deleted.  Inactive if PKA and Bcy1 is present at the same time as Krh is active or cAMP or PKA is absent.(T. Peeters et al. 2006; Takashi Toda et al. 1987; T Toda et al. 1987) |
| Rim15 | Present unless deleted | Always unphosphorylated unless PKA is active and Rim15 is present. (Swinnen et al. 2006).  Phosphorylated if Rim15 is present and Sch9 is phosphorylated. (Wanke et al. 2008). | - |
| Gis1 | Present unless deleted | Always unphosphorylated unless Rim15 is unphosphorylated and Gis1 is present.(Swinnen et al. 2006). | - |
| Msn2,4 | Present unless deleted | Always unphosphorylated unless Rim15 is unphosphorylated and Msn2,4 is present.(Swinnen et al. 2006). | - |

Table 3. Rules and references associated to any field of the Boolean vector in the TOR pathway.

| TOR | | | |
| --- | --- | --- | --- |
| Name | Presence | Phosphorylation | Specific activity |
| EGO | Present unless deleted | - | Always inactive unless NH3 and EGO is present. (Binda et al. 2009; Bonfils et al. 2012; Bar-Peled et al. 2013) |
| Tor1 | Present unless deleted | - | - |
| Tor2 | Present unless deleted | - | - |
| Kog1 | Present unless deleted | - | - |
| Tco89 | Present unless deleted | - | - |
| Lst8 | Present unless deleted | - | - |
| TORC1 | Always absent unless if Tor1 or Tor2 is present as well as Kog1, Toc89 and Lst8. (Reinke et al. 2004) | - | Always inactive unless EGO is active and TORC1 is present. (Binda et al. 2009)  Inactive if Snf1 is phosphorylated and EGO is active and TORC1 is present. (Hughes Hallett, Luo, and Capaldi 2014). |
| Sch9 | Present unless deleted | Always unphosphorylated unless TORC1 is active and Sch9 is present. (Urban et al. 2007) | - |
| Sfp1 | Present unless deleted | Always unphosphorylated unless TORC1 is active and Sfp1 is present. (Marion et al. 2004; Lempiäinen et al. 2009). | - |
| Tap42 | Present unless deleted | Phosphorylated when TORC1 is active and Tap42 is present.  Unphosphorylated if Tap42 is present and TORC1 is inactive or if Tap42 is absent. (Beck and Hall 1999). (Jiang and Broach 1999; Di Como and Arndt 1996). (Yan, Shen, and Jiang 2006).  Phosphorylated when Snf1 is phosphorylated and Tap42 is present. (Hughes Hallett, Luo, and Capaldi 2014).  Unphosphorylated if Tap42 is present, TORC1 is inactive and Snf1 is unphosphorylated. (Hughes Hallett, Luo, and Capaldi 2014). | - |
| PP2A | Present unless deleted | - | Always active unless Tap42 is phosphorylated and present and PPA2 is present or if PPA2 is absent. (Beck and Hall 1999). (Jiang and Broach 1999; Di Como and Arndt 1996). (Yan, Shen, and Jiang 2006). |
| Mks1 | Present unless deleted | Phosphorylated if TORC1 is active and Mks1 is present, or if GLUex is absent and Mks1 is present. (Hughes Hallett, Luo, and Capaldi 2014).  Unphosphorylated if Mks1 and PP2A is present and PP2A is active or if Mks1 is absent. (Dilova et al. 2004; Broach 2012). | - |
| Rtg2 | Present unless deleted | - | - |
| Rtg1,3 | Present unless deleted | Always phosphorylated unless Mks1, Rtg2 and Rtg1,3 is present and Mks1 is unphosphorylated. (Dilova et al. 2004; Broach 2012). | - |
| Gln3 | Present unless deleted | Always phosphorylated unless PP2A is present and phosphorylated and Gln3 is present. (Georis et al. 2009; Broach 2012; Conrad et al. 2014). | - |
| Gat1 | Present unless deleted | Always phosphorylated unless PP2A is present and phosphorylated and Gat1 is present. (Georis et al. 2009; Broach 2012; Conrad et al. 2014). | - |

Table 4. Rules and references associated to any field of the Boolean vector for the enzymes not specifically assigned to a pathway.

| Enzymes | | |
| --- | --- | --- |
| Name | Presence | Phosphorylation |
| PFK2 | Present unless deleted | Always unphosphorylated unless PKA is active and PFK2 is present. (Dihazi, Kessler, and Eschrich 2003) (Active when phosphorylated.) |
| TREH | Present unless deleted | Always unphosphorylated unless PKA is active and TREH is present.(Schepers et al. 2012) |
| PK | Present unless deleted | Always unphosphorylated unless PKA is active and PK is present. (Portela et al. 2002) |
| FBP1 | Present unless deleted. Present is FBP1 is active as target. | Always unphosphorylated unless PKA is active and FBP1 is present. (Rittenhouse, Moberly, and Marcus 1987). |
| ACC | Present unless deleted | Always unphosphorylated unless Snf1 is active and Snf4 and ACC is present as well as any of Sip2, Sip1 or Gal83.(Woods et al. 1994) |
| HXK2 | Present unless deleted | Always unphosphorylated unless HXK2 is present and glucose is not present. (Fernández-García et al. 2012) |

Table 5. Rules and references associated to any field of the Boolean vector for metabolites.

| Metabolites | |
| --- | --- |
| Name | Presence |
| GLUex | 1 |
| ATP | Always present unless deleted. |
| cAMP | Present when ATP is present and AC is active.  Absent when AC is inactive and PDE is present and phosphorylated. (Sass et al. 1986; Nikawa, Sass, and Wigler 1987; Ma et al. 1999; Hu et al. 2010) |
| F16BP | Always absent unless GLUex is present. |
| NH3 | 1 |

Table 6. Rules and references associated to any field of the Boolean vector for the gene targets or target groups of the transcription factors.

| Targets | |
| --- | --- |
| Name | Activity |
| PDS | Always inactive unless Gis1 is phosphorylated. (I Pedruzzi et al. 2000; Martínez-Pastor et al. 1996). |
| STRE | Always inactive unless Msn2,4 is phosphorylated. (I Pedruzzi et al. 2000; Martínez-Pastor et al. 1996). |
| RTG | Always inactive unless Rtg1,3 is unphosphorylated and present. (Dilova et al. 2004; Broach 2012; Liu and Butow 1999) |
| NCR | Always inactive unless Gat1 is present but not phosphorylated or is Gln3 is present but not phosphorylated. (Georis et al. 2009; Broach 2012; Conrad et al. 2014). |
| SUC2 | Always inactive unless Mig1 is phosphorylated(Westholm et al. 2008; Santangelo 2006; Broach 2012; Schüller 2003). |
| ADH2 | Inactive if Adr1 is phosphorylated and PKA is active. Always inactive unless Adr1 is phosphorylated. (Turcotte et al. 2010; Soontorngun et al. 2012; Broach 2012; Kacherovsky et al. 2008; Smith et al. 2011; Cherry et al. 1989) |
| CSRE | Always inactive unless Sip1 or Cat8 is phosphorylated. (Broach 2012; Leverentz and Reece 2006; Turcotte et al. 2010; MacPherson, Larochelle, and Turcotte 2006). |
| PCK1 | Always inactive unless Sip1 or Cat8 is phosphorylated. (Broach 2012; Leverentz and Reece 2006; Turcotte et al. 2010; MacPherson, Larochelle, and Turcotte 2006). |
| FBP1 | Always inactive unless Sip1 or Cat8 is phosphorylated. (Broach 2012; Leverentz and Reece 2006; Turcotte et al. 2010; MacPherson, Larochelle, and Turcotte 2006). |
| Ribosomal | Always inactive unless Sfp1 is phosphorylated. (Marion et al. 2004; Lempiäinen et al. 2009). |

#### 1.5 Dynamics of the Boolean model indicate either a model gap or rate differences in the pathways.

When implementing the crosstalk, we found that the dynamics did not operate according to literature in contrast to the steady state result. In contrast to literature, Adr1 was inactivated by PKA instead of Snf1(Cherry et al. 1989) and PKA acted as the main regulator of Rim15 instead of Sch9 (Ivo Pedruzzi et al. 2003). This could either indicate that these pathways may not operate on the same time scale or the complexity of the pathways are not equally known. When iterating over discrete time steps, one does not consider time but complexity of the modelled pathways. For instance, if a pathway is well described and can be model in great detail, it takes many iterations to reach a steady state. In contrast, a poorly understood pathway may need very few iterations to reach the steady state although in reality, signaling via the poorly understood pathway may take more steps than via the well-annotated pathway. In this work, a synchronous modelling scheme was used, meaning that at each iteration the state vectors are updated simultaneously as there is little information on order and duration of state transitions available (Garg et al. 2008). In these cases, it is hard to say if the simulations reflect the reality, indicates that the PKA pathway is less understood than the Snf1 pathway or if the discrepancy is a result of difference in rates between the pathways. Either way, these results highlight the lack of understanding in the dynamics of signal transduction in nutrient signaling pathways.

#### 1.6 Knock out of major signaling components reveals question marks about reported crosstalk mechanisms.

In the deletion experiments in the Boolean model during high nutrient availability, the simulation of Reg1 knockout showed almost the same effect on the SNF1 pathway as nutrient depletion. Only Adr1 activity was not affected which opposes the observations by Dombek et al. (1999) that described constitutive ADH2 expression in Reg1 mutant cells (Dombek et al. 1999). Since Adr1 is the main regulator of ADH2 activity, the previously discussed inhibition of Adr1 activity by active PKA (Cherry et al. 1989) may be the reason for these contradicting results and thus, the relevance of this crosstalk may be questioned.

In literature the Snf1 knock out is described to have a phenotype resembling over activation of PKA, however in our simulated deletion experiments in the Boolean model during low nutrient availability, AC was activated but the Krh proteins inhibit PKA if no glucose is present. PKA activation is a fine-tuned process that requires more complexity such as high cAMP concentrations upon Krh activation (T. Peeters et al., 2006) which could not be modelled using our Boolean approach.

### 2 Hybrid model

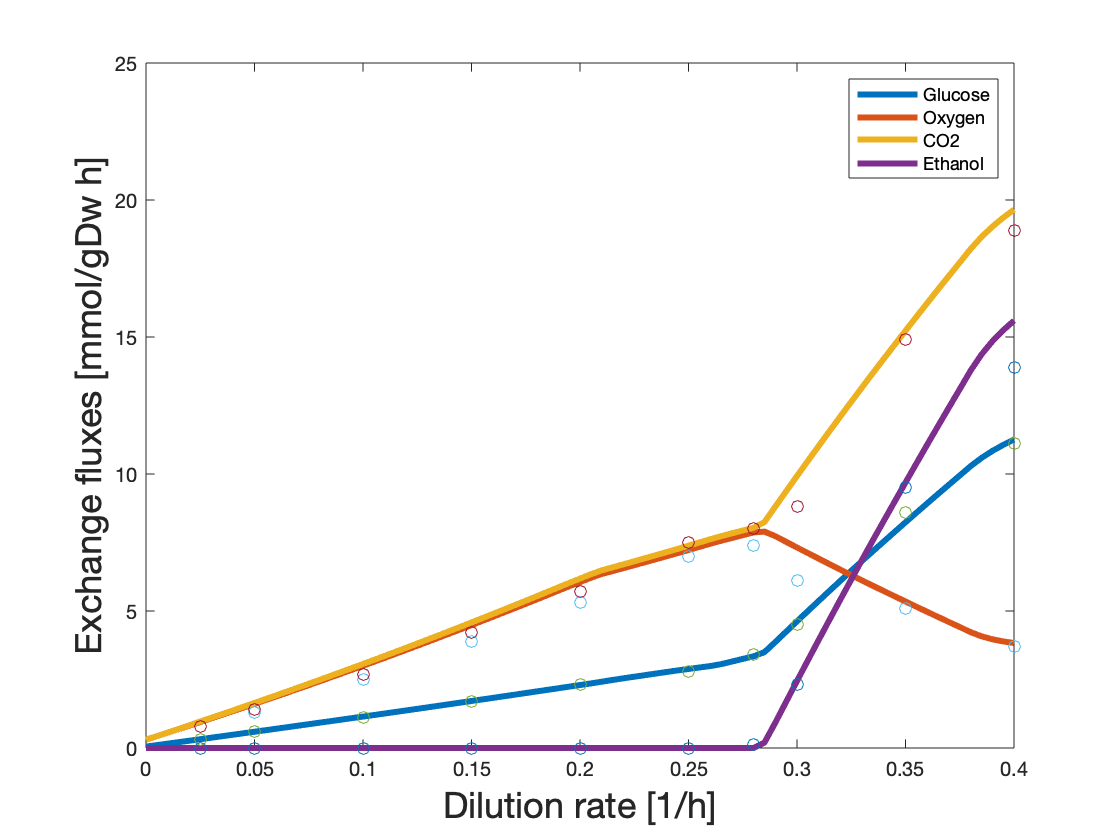

Supplementary Figure 3. Exchange fluxes for the hybrid model plotted over experimental data. Simulations showed a median relative error of 9.82% in the whole range of dilution rates from 0 to 0.4 h^-1^.

Supplementary Table 7. Summery of the statistics done comparing the ecModel and the hybrid model in their ability to predict protein abundance.

| Statistics | Respiration | | Fermentation | |
| --- | --- | --- | --- | --- |
|  | ecModel | Hybrid model | ecModel | Hybrid model |
| PCC (statistics) | 0.23379 | 0.18143 | 0.013517 | -0.038213 |
| PCC (P-value) | 0.019 | 0.048 | 0.417 | 0.648 |
| KS (statistics) | 0.50575 | 0.41379 | 0.36667 | 0.0008511 |
| KS (P-value) | 1.7062e-10 | 3.636e-07 | 1.1208e-07 | 0.25 |
| r | 2.62 | 1.56 | 3.56 | 2.33 |

As can be seen by the summery statistics, prediction of individual proteins are difficult. Yet, the ecModel predicts 48.3% of the proteins within one order of magnitude in the respiration state and 32.5% of the proteins in the fermentation state and for the hybrid model this is 65.5% and 40.8% respectively. We defined miss-predicted proteins as deviating with over one order of magnitude and divided them into two groups, the moderately miss-predicted proteins that were over- or underpredicted with 1-5 orders of magnitude and the heavily miss-predicted proteins that were miss-predicted with over 5 orders of magnitude. In the latter group we have only proteins that are either not detected in the experimental data but used by the model or proteins not used by the model but detected by experimental data, in the case of the hybrid model. This can largely be explained by the model having a preferential use of isoenzymes and that membrane proteins are often not detected in mass spectrometry studies. However, there are a few exceptions of some proteins lying in the pathways that makes out the precursors for the biomass. By adding the signaling layer to the ecModel we force the use of isoenzymes and pathways that might not be used in that particular steady state which decreases the number in the category of heavily miss predicted proteins from 32.2% to 13.7% in the respiration state and from 35.0% to 17.5% in the fermentation state. In the moderately miss predicted protein category, the proteins were on average miss predicted by 1.80 orders of magnitude by the ecModel in the respiration state and by 1.71 orders of magnitude by the hybrid model. In the fermentation state the ecModel were off by on average 1.49 orders of magnitude whereas the hybrid model was off by 1.61 for the moderately miss predicted proteins.

One of the contributing factors to the miss-predicted proteins could be the kcat curating process. The kcats are curated trough GECKO; in case the kcats are not available for *Saccharomyces cerevisiae* the kcats are taken from the phylogenetically closest related organism that the value does exist for. This process will make sure we have some constraints based on enzyme kinetics but will also contribute to errors in our predictions where the values differ between organisms. Predicting proteins based on optimality principles has the inherent problem that when confronted with several choices of how to carry a flux trough a reaction, the determining factor will lie heavily on the molecular weight of the proteins involved in the reaction and the flux needed based on the steady state assumption. In reality cells need to be prepared for changes in the environment and are seldom in a steady state, meaning they also need to put energy into enzymes not needed for the specific conditions. We also see a trend in both respiration and fermentation that the glycolysis, TCA and PPP proteins are overpredicted while the OPP proteins are underpredicted. This might be due to two factors, the glycolysis, TCA and PPP mainly consists of globular proteins and the OPP proteins are mainly membrane bound. There is an intrinsic difficulty in mass spectrometry studies to quantify membrane proteins which might contribute to the appearance of a higher relative abundance of globular proteins compared to membrane bound proteins.

#### 2.1 Multiscale modeling allows the discovery of complex interactions.

When running the individual models, we gain different insights. When running the Boolean model we can replicate the steady state results from literature and we get indications of knowledge gaps we have when it comes to the dynamics of the signaling transduction pathways and the crosstalk between the pathways. When running the ecModel we get a good fit to the exchange rates, but when adding regulation, we can also improve prediction power of the individual enzymes in the model. From the Boolean model, we can see that in respiration condition Gis1 mainly acts as activator of glycerol metabolism, Msn2/4 is activation STRE elements, Adr1 activates the genes needed for utilization of non-glucose carbon sources and Sip4 and Cat8 activates promoters with carbon source response elements (CSREs). From just the Boolean model it is hard to predict the findings we got when combining the model, especially the connection between Snf1 and chronological lifespan.

#### 2.2 Robustness is reflected by the basal expression of pathways not used in steady state

In the hybrid model we get futile flux trough reactions not used by the model which reflects that the regulation forces to produce enzymes are not needed for the objective function. We argue that this adds a robustness to the cells since natively they are seldom in steady state and thus a basal expression is needed for the robustness and fitness for survival. However, that comes at a cost of not utilizing the enzymes in an optimal way. As an example, we can look at the galactose pathway in respiration. GAL10 and PGM2 have a base expression but are not used to produce biomass, thus only carry futile flux. PGM2 is induced by galactose and repressed by glucose (Oh and Hopper 1990) however, when looking at the fit to the proteomics data we have a good prediction on the PGM2 protein. This can be observed for other carbon source reactions such as ADH1.

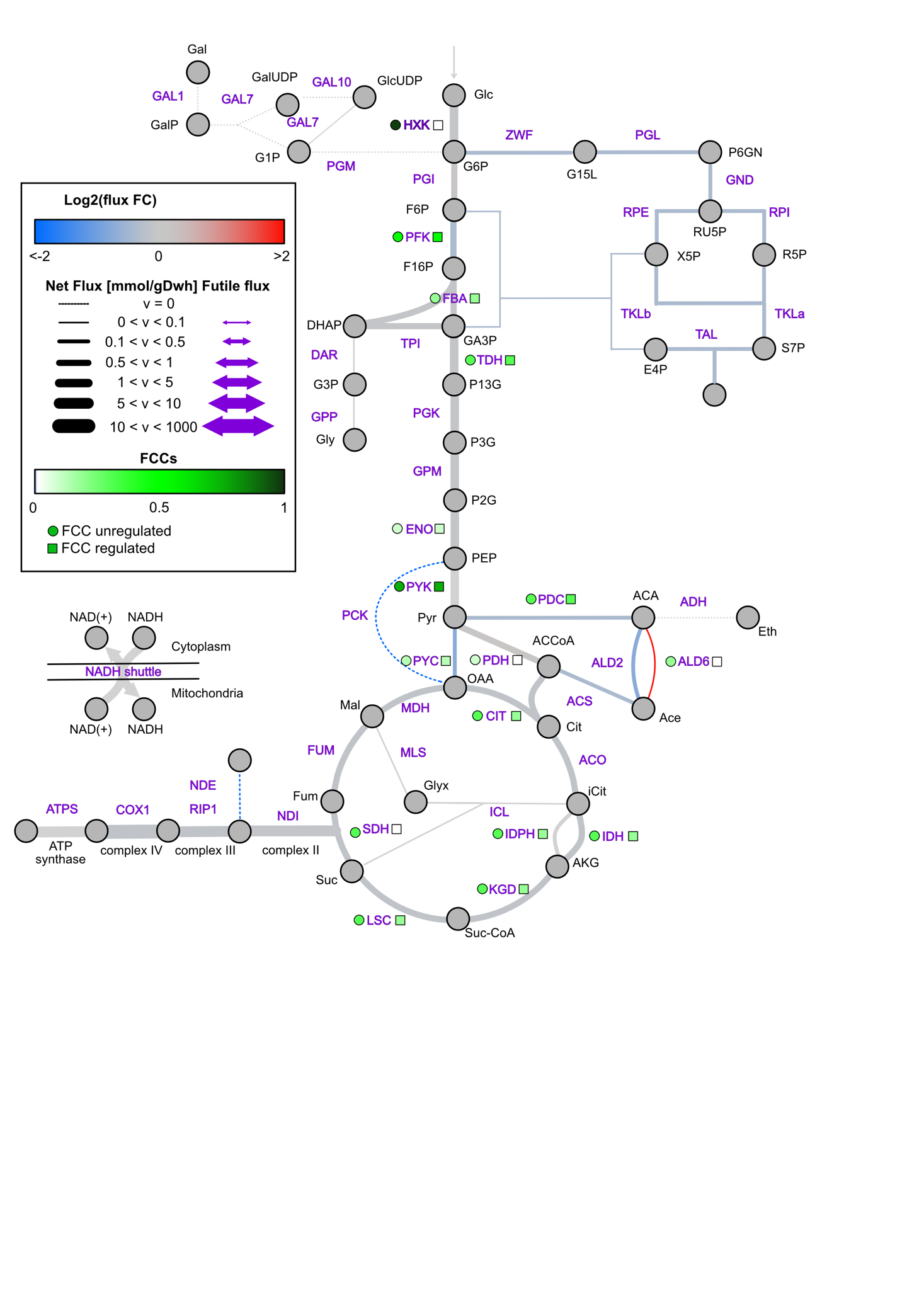

Supplementary Figure 4. The fluxes trough the core reactions in the metabolism are represented by the width of the connectors where dotted lines represent zero flux. The colour of the connectors represent the change in flux from the wilde type (WT) hybrid model compared to the SNF1 deletion hybrid model. The FCCs are represented in the model where the WT are compared to case with SNF1 deletion case.
